# Rubin-vase percept is predicted by prestimulus coupling of category-sensitive occipital regions with frontal cortex

**DOI:** 10.1101/107714

## Abstract

Fluctuations in what an observer perceives can be demonstrated in images that give rise to competing percepts, examples include binocular rivalry, figure-ground illusions, and other phenomena. These stimuli are an important and well-studied stimulus set in studying human conscious perception.

One such stimulus is the Rubin’s vase illusion, a bistable stimulus that gives rise to the perception of a vase, or the profile of two faces (hereafter referred to as the face-vase stimulus). Research using this stimulus has predominately focused on the neural response whilst viewing it (Andrews et al., 2002; Hasson et al., 2001; Kleinschmidt et al., 1998). An explanation for the phenomenological switch could exist in distinct features of the on-going brain activity, Hesselmann, Kell, Eger, & Kleinschmidt (2008) showed that fluctuating BOLD levels in Fusiform Face Area (FFA) prior to presentation of the face-vase stimulus bias the participants’ perception. These findings corroborate earlier reports of binocular rivalry (Tong et al., 1998) and face-vase illusions (Andrews et al., 2002; Hasson et al., 2001; Kleinschmidt et al., 1998), wherein enhanced BOLD in FFA during face percept was reported. Enhanced prestimulus neural activity in category variant nodes may provide a predisposition of the biasing competition. Prestimulus periods can impact auditory perception (Sadaghiani et al., 2009), predict perceptual biases (Bode et al., 2012), and basic visual perception (Busch et al., 2009). M/EEG methods are ideally suited to investigate the patterns of prestimulus brain states that predict upcoming perceptions, due to the strength in temporal resolution, and ability to look at large scale network dynamics. Nonetheless, research on bistable images reports utilizing M/EEG has been rare (Britz et al., 2009; Doesburg et al., 2009).

There is strong evidence that conscious experience requires large-scale activation of prefontal-parietal networks during the post-stimulus period. The evidence in this period is best demonstrated utilising the contrasts of masked vs unmasked words or detected sounds vs non-detected sounds (Dehaene and Changeux, 2011). However, there is strong evidence of prestimulus activity influencing conscious awareness (Ruhnau et al., 2014). Here we test the windows to consciousness (Win2Con) framework of consciousness, a framework outlined by Ruhnau et al. (2014) and Weisz et al. (2014). The framework was initially derived from the near-threshold (NT) example (wherein subjects report either seeing or not seeing the stimulus), and was driven to test directly these conscious awareness states. The hypothesis of the framework runs as follows: Conscious awareness requires that stimulus variant nodes (i.e. areas that have specialised functioning, e.g. the visual cortex) propagate a representation to higher-order neural areas (i.e. frontal-parietal) in agreement with Dehaene & Changeux (2011). How this propagation occurs in the post-stimulus period can be debated, but the framework makes specific claims that this propagation is afforded by an open state of communication before stimulus onset. As a prerequisite for conscious access to NT stimuli this hypothesis has had supportive findings (Leske et al., 2015; Weisz et al., 2014). Unlike the NT contrast in conscious processing, the intriguing aspect about bistable images is that on each occasion an object specific conscious content is associated with the stimulus. Herein we wish to test the framework with conscious content and see if this prerequisite of an open state of communication holds true for the case of second-order visual items (i.e. houses and objects etc.). With this knowledge we can redefine the framework with the presented empirical evidence.

To measure the contribution of the essential nodes we look at the levels of local excitability two-fold via oscillatory activity (alpha to low beta), and evoked response. Alpha/low-beta oscillatory activity putatively reflects the excitability of brain regions (Haegens et al., 2011). Strong power being related to inhibition of brain regions resulting in decreased stimulus processing and vice versa for low levels of power (Jensen and Mazaheri, 2010). Likewise, the evoked response can highlight task specific essential nodes, and although they can only be tested in the post-stimulus period we will use the evoked response to inform the analysis. With the regional information of the essential nodes uncovered, we can then test the open window of communication in the prestimulus period using a functional connectivity measure. In the face-vase stimulus the notion is that the two contents of the stimulus will elicit two two essential nodes (i.e. face processing node, and object(vase) processing node). What is then crucial for the Win2Con framework is how the network states of the two essential nodes predict the upcoming conscious content, with a prediction that they will bifurcate.

## Materials & Methods

### Participants

20 right-handed volunteers with normal or corrected-to-normal vision participated in this experiment (9 m/11 f, mean age 25.3). During the course of the experiment participants wore non-magnetic clothes, and a questionnaire prior to the experiment excluded any metal artefacts on the participants being. The Ethics Committee of the University of Trento approved the experimental procedure and all participants gave written informed consent before taking part in the study. Two participants had to be excluded due to an excessive amount of artefacts.

### Experimental Procedure

After the placement of the Head Position Indicator (HPI) coils to the participants’ head, the experiment proper began. Participants were seated upright in the MEG system. They were instructed to keep fixation, and to avoid eye blinks and movements as best as possible during the experiment. In between the blocks participants had a short break but remained seated in the MEG system. Visual stimuli were displayed via a video projector outside of the MEG chamber and projected to a back-projection screen in the MEG chamber. A camera allowed the investigator to monitor participants during the experiment.

During a trial a fixation cross would appear at the centre of the screen, this would last between 1-1.8 sec. After this jittered period the face-vase picture was displayed at the centre of the screen for 150 msec (see Figure 1 for trial example). Half of the participants were presented the original face-vase picture (black background) and the other half presented with a colour inverted face-vase picture (white background). This was done to ensure that the luminance of the picture did not bias the dominant percept, and post-hoc group analysis revealed no differences between the differing background on the measures reported. After the face-vase picture presentation a mask stimulus was presented for 200 msec. Scrambling blocks of pixels of the face-vase picture randomly created this mask (see Figure 1). After the presentation of the mask participants were asked to respond if they saw the face or the vase. The response window was 2 seconds and if participants did not respond then the experiment would continue. Participants responded with the index and middle finger of the left or right hand. The response hands were counterbalanced over subjects. If participants experienced both precepts during a trial, they were instructed to report the first percept experienced. There were a total of 400 trials, broken into 4 blocks of 100 trials. The presentation of visual stimulus material during MEG recordings was controlled using Matlab and the Psychophysics Toolbox (Pelli, 1997), with timings corrected using a photo diode. The procedure of the experiment is illustrated in Figure 1.

**Figure 1:**
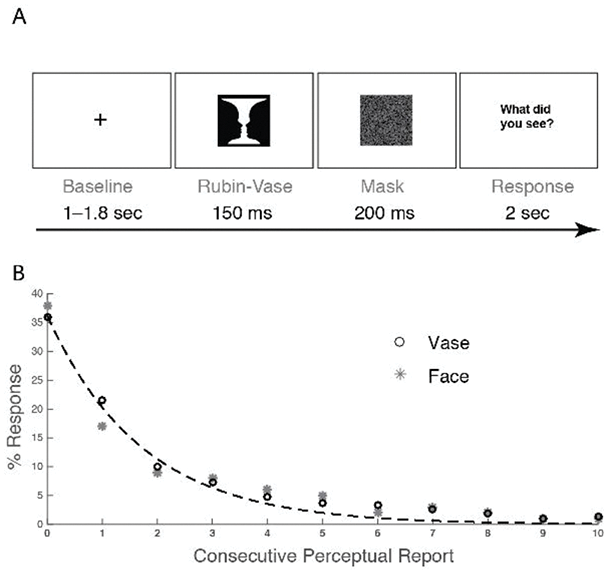
A: Experimental Trial. B: The distribution of responses based on the amount of repetitions of percept response was reported (goodness-of-fit R^2^ = 0.96 for face, R^2^ = 0.98 for vase). For example, overall participants would say face twice in a row on 17% of trials. Dashed line indicates the binominal distribution.

### Behavioural Analysis

We collected behavioural reports after the end of each trial, giving us 400 responses. To test for the stochastic nature of the response we used curve-fitting procedures from the Matlab (The MathWorks, Natick, MA, Version 8.2.0 R 2013b) curve-fitting toolbox. Specifically, for each participant we binned the data according to how many trials in a row they responded with the same perceptual report. We broke this down in 10 bins with 0 repetition to 10 repetitions, this data was then fitted to a binomial distribution generated in Matlab across the 10 bins. The two distributions were then fit using a goodness-of-fit measure. We also ran a drift diffusion model (DDM) analysis on the data. We wanted to test if there was any choice history priming. The analysis ran as follows: First, we collapsed across all participants and coded 1 as vase and 0 as face perception (as there is no erroneous response); Second, we classified the data into two conditions, one where the preceding trial was different report, and the second where the preceding trial was the same report. Finally, we tested the DDM using the DMAT toolbox (Vandekerckhove and Tuerlinckx, 2008), and tested two models; one which tested the null effect and the other which tested the starting bias effect. The difference between the two models deviance was the statistical analysis used to ascertain which model best fitted the data.

### MEG Data Acquisition

The MEG recordings were carried out using a 306-channel whole-head VectorView MEG system (Elekta-Neuromag, Ltd., Helsinki, Finland, 204 gradio- and 102 magnetometers) installed in a magnetically shielded chamber (AK3b, Vakuumschmelze Hanau, Germany), with signals recorded at 1000Hz sample rate. Hardware filters were adjusted to band-pass the MEG signal in the frequency range of 0.01Hz to 330Hz. Prior to the recording points on the participant's head were recorded using a digitizer (Polhemus, VT, USA). These points included the 5 HPI coils, the three fiducials (nasion, left and right preauricular point), and over 200 additional points on the head. The HPI coils were used to monitor the head position during the experiment.

### MEG Preprocessing

From the raw continuous data, we extracted epochs of 4 seconds lasting from 2.5 seconds before onset of the picture to 1.5 seconds after onset of the picture. This resulted in 400 trials per participant. This epoched data was first high-pass filtered at 1 Hz (IIR Butterworth 6-order two-pass filter with 36 dB/oct roll-off), followed by bandstop filter of 49 – 51Hz to remove power line noise. Trials were visually inspected for artefacts (muscle artefacts, eye blinks, channel jumps) and the contaminated trials rejected. For each participant, the trials were then assigned to the 2 conditions according to the participants’ response. To ensure a similar signal-to-noise-ratio across conditions, the trial numbers were equalized for the compared conditions by random omission. Meaning that in the resultant data the percentage of trials left in the analysis was: M = 79.32%, SD = 15.12. Data was down sampled to 400 Hz using the resample function of Matlab and then was subsequently detrended using a single order polynomial (i.e. linear detrend).

### Common Source Method

Sensor level data was projected into source space using a whole brain equally spaced grid linear constrained mean variance (LCMV; (Van Veen et al., 1997)) beamformer approach. For all source reconstructions an anatomically realistic head-model (Nolte, 2003) was created using individuals structural MRI (15 out of the final 18 participants) and the Polhemus derived scalp shape. A three-dimensional source grid (resolution: 1.5 cm) covering the entire brain volume was calculated using both sensor types (magnetometers and gradiometers). For each one of these points we constructed a virtual sensor, essentially upsizing the approach commonly used for one virtual sensor (http://www.fieldtriptoolbox.org/tutorial/shared/virtual_sensors) to the 889 points in the source grid. To obtain this we first generated common spatial filter weights (i.e. using all trials, using specific time period of the covariance window). These spatial filter weights were then multiplied by the raw data of the two trial types. This raw time series then underwent the specified analysis (e.g. time frequency analysis). It is then interpolated onto individual MRIs for 15 participants, while for three participants without MRI scan we used a template MRI which was morphed to fit the individuals head shape using an affine transformation. The interpolated activation volumes were then normalized to a template MNI brain provided by the SPM8 toolbox (http://www.fil.ion.ucl.ac.uk/spm/software/spm8). It is then these activation volumes that form the base of the statistical test that follows.

### Prestimulus Spectral Power Analysis

Task-related changes in oscillatory power were estimated from 1 to 29 Hz in steps of 1 Hz using a sliding window FFT time-frequency transformation. Time windows were adjusted per frequency (time window: Δt=4/f sliding in 50 msec steps) and Hanning-tapered. Power was calculated for 102 magnetometers and 102 combined planar gradiometers (the latter resulting from 204 gradiometers that were summed to a single positive-valued number at each sensor). Based on visual inspection of the uncorrected sensor results between 1-29Hz the frequency range was limited to 5–20 Hz to increase the sensitivity of the non-parametric testing. In order to test if power modulations are significantly different between conditions (face and vase percept) we performed a nonparametric cluster-based permutation t-test (Maris and Oostenveld, 2007) on the time-frequency representations of the two conditions. The dimension of the inputted data were 21 time points (-1000 to 0 msec prior to stimulus onset in 50 msec intervals) by 16 frequencies (5-20Hz). For the non-parametric cluster selection, we used the maximum sum t-stat and this was tested against 1000 randomisations. This procedure was implemented for each sensor type separately. Spatial neighbours were defined as sensors with a maximal distance of 4 cm, resulting in an average of 4.6 neighbours per channel. For the source analysis we followed the above described procedures using a time period of −1000 to −200 msec before stimulus as the covariance window. Near identical input parameters of statistical analysis used in the prestimulus sensor analysis were used, with the only difference being the spatial neighbourhood now being composed of voxels, there was an average of 7.4 neighbours per voxel (maximal distance = 1.6 cm). The resultant statistics are corrected for multiple comparisons using the non-parametric cluster based approach (Maris and Oostenveld, 2007).

### Event-related fields analysis

For the analysis of the event-related fields we first bandpass-filtered the data between 1 and 25Hz and subtracted a −200msec to 0msec baseline. We then applied a dependent-samples *t*-test with the non-parametric Monte-Carlo correction (see above), using the threshold parameters of p < .05 and the maximum sum of the cluster as the cluster statistics. A cluster had to include a minimum number of at least 2 sensors. The distribution of our statistics was estimated from 1000 randomisations. This test was performed across a window of 100 to 500msec locked to the stimulus presentation. The contrast again being between the trials where participants reported a vase vs. reporting a face. The tests for the magnetometers and the combined planar gradiometers were done separately. The statistical outcome of this analysis was used to inform the analysis approach in source space. Again we used the LCMV taking a covariance window of 250msec to 450msec. Source-level time series were averaged and the absolute value was computed. Changes to baseline (-200 to 0msec) were expressed as relative changes in order to counteract the depth bias common in beamforming. Taking the significant time period, we selected the maximal peak t value and used that as the most likely generator, as when applying non-parametric correction no significant voxels/time-points were present.

### Seeded Connectivity

For the analysis of functional connectivity we used the imaginary part of coherency (iCOH), a connectivity measure insensitive to volume conduction (Nolte et al., 2004). Temporally, we focussed the analysis on the time window given by the prestimulus power (−1 to −0.6 s) and across the 7 – 15 Hz range, this range was selected as visual inspection of the iCOH between 5-20Hz showed a peak effect in this frequency range. The shorter time window was selected due to the period of significant effects in the prestimulus source effects. Using the aforementioned source approach and a −1 to −0.6 sec covariance window we generated a grid to grid point iCOH from the Fourier spectrum between 7 and 15 Hz. 1 grid point was then selected as the seed for the iCOH measure giving us a source representation of the seed x source x frequency connectivity. These source maps of iCOH were calculated separately for the Vase and Faces perceptual report using the two ROIs highlighted in previous analysis (the LO, and OFA) as seeds. For the seeded connectivity we looked at the interaction of the 2 (Seed: LO and OFA) by 2 (Report: Vase and Face) factors. This was calculated using a dependent sample t-test to look at the difference of the contrasts between the OFA Seed (i.e. Face Report OFA Seed Minus Vase Report OFA Seed) and the LO seed (i.e. Face Report LO Seed Minus Vase Report LO Seed). This was done across the 7 to 15 Hz frequency range, more explicitly the interaction test is a first omnibus approach to identify relevant brain regions that show differential connectivity effects for OFA and LO seeds with respect to the actual perceptual report (see Figure 3). To illustrate how this interaction is driven, iCOH values from the respective seed x report combination were extracted from the region exhibiting the significant cluster level interaction effect, these values were corrected using false discovery rate multiple comparison correction.

## Results

The current study aimed at disentangling prestimulus brain activity determining the dominant percept of an ambiguous picture, here the face-vase stimulus. We investigated brain activity on a local and on a network level in the time interval before the picture was shown and with a focus on low-frequency oscillatory power. As only one essential node was present in the prestimulus period we also analysed the window after stimulus onset, and focused here on the evoked response.

### Behavioural

The report of a vase or face was as equal as likely overall (Mean: 49.67 %, SD: 13.10%, across participant range 22% to 84%, with a t-test against chance (50%) showing non-significance *t (17)* = 0.11, p = .92). The reaction times were not significantly different *t (17)* = −0.37, p = .72, (vase M = 619msec SE = 44; face M = 631msec SE = 42). As in Hesselmann et al. (2008) we wanted to ascertain if the reported perception was stochastic trial-by-trial. As such, we performed analysis on the sequences of reported percepts binning the trials into a range of 0 to 10 repetitions, and tested this against a binomial test. For both the vase and the face a binomial distribution was shown (goodness-of-fit R^2^ = 0.96 for face, R^2^ = 0.98 for vase) indicative of no systematic reporting. That is, during each trial a participant was equally likely to report a face or vase irrelevant of the previous trial. Finally, the DDM outcome highlighted that there was a better fitting model when taking into account a starting point bias, with the repetition showing a bias and the no repeat not showing a bias (χ^2^Diff = 274.28, p<0.001). It would seem that the brief presentation and masking elicited our required effect that there was an unstable percept occurring trial-by-trial and there were no carry-over effects from the previous trial. Although reporting the same stimulus one trial after reporting the same did elicit effects of the starting bias within the drift diffusion model.

### Prestimulus Effect

In a first step, we assessed prestimulus low-frequency power at sensor level and compared activity preceding the perception of the two faces versus the vase. We did that for both sensor types separately. Planar gradiometers showed a significant power increase (cluster p= 0.02, 8–20 Hz, 1 sec to 200 msec before picture onset) encompassing left frontal and right posterior sensors for subsequent reported faces. The magnetometers showed no significant differences between reports (max cluster p = 0.09, 11–20 Hz, 1 sec to 400 msec before picture onset, with a mainly right posterior topography; See Inline Supplementary Figure 1). In both cases the frequency showing a continuous effect over the whole time window was the 12-17 Hz range.

In order to get a better estimate of where in the brain the low-frequency power differences derive from, we performed an LCMV virtual-sensor source analysis of the power modulations observed at sensor level (12–17 Hz, −1– −.2 sec). The source analysis showed one cluster of significant difference in the 12-17Hz range which covered the time period 1 sec to 600 msec before stimulus onset. Source results indicate a spatial cluster that encompassed a region of the right Lateral Occipital Complex (LO; MNI *x* = 44 *y* = −71 *z* = 12) and extended into the temporal sulcus (p = .02, *corrected*, see Figure 2A).

**Figure 2:**
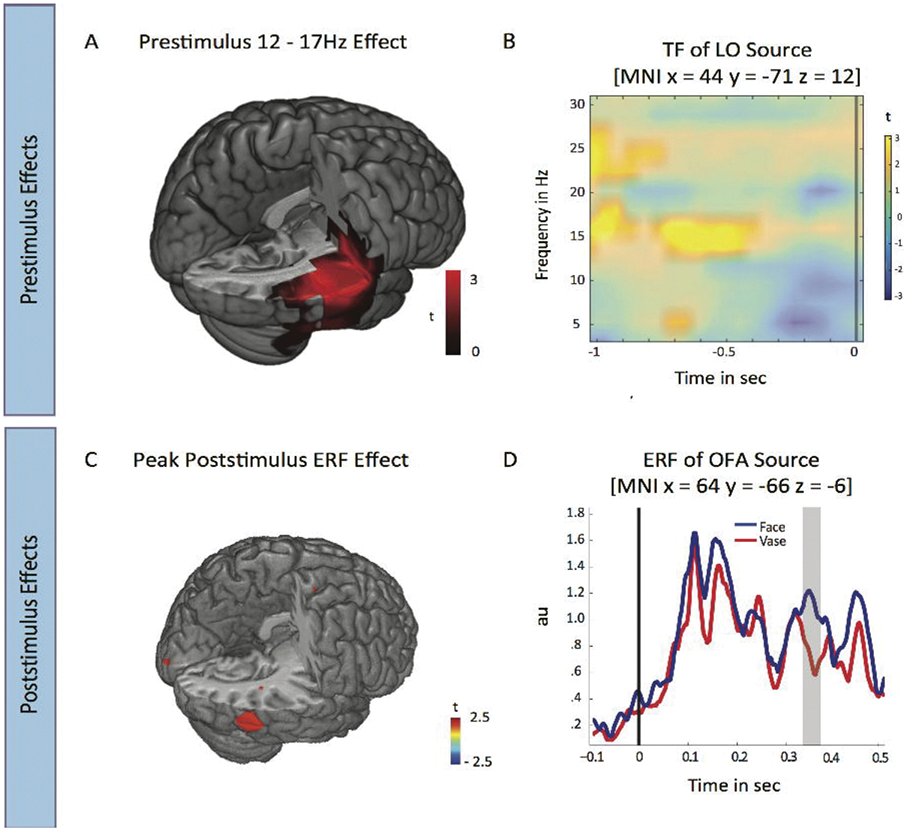
A: A brain cut-out corrected statistical map of the t score differences between the two reports (12–17 Hz, 1 to 0.2 sec prestimulus) with positive values indicating more power for face report vs. vase report. Power is significantly increased prior to face reports in the right LO (MNI *x* = 44 *y* = −71 *z* = 12), in addition to left frontal sources (not shown here). The map is corrected for multiple comparisons and shows colour where significance passes the boundary of this correction. B: Source TF of the selected power effect in LO. Increased power prior to Face reports can be seen in particular ~15Hz. Black line denotes stimulus onset. Masked at p < .01. This was reconstructed from an LCMV source reconstruction from the same right LO using a virtual sensor approach. C: shows the source reconstructions with an LCMV beamformer averaged on the 350msec point; notice the OFA (MNI *x* = 64 *y* = −66 *z* = −6) locus of the highest stat (p < .01, *uncorrected).* D: shows the evoked response source waveforms for the two conditions using the virtual sensor at the centre of the OFA region. Black line denotes stimulus onset, and grey shaded area the window of significance in the sensor space. Values are arbitrary units of evoked response.

### Event-related fields and sources

The second analysis looked at the event related fields time-locked to the stimulus onset, and contrasted the two perceptual reports. Using a 100 to 500 msec window, the combined planar gradiometers showed a significant difference between face and house reports at the time period 341 and 375msec after the trial presentation (p = .019). The gradiometer topography shows a right temporal and bilateral occipital morphology (See Inline Supplementary Figure 2). Source projection of this significant time window indicating a maximal peak at the 350msec time period within the significant difference being spatial located in the right Occipital Face Area (OFA; MNI *x* = 64 *y* = −66 *z* = −6; p <. 01, un*corrected*, Figure 2C), a region of that is consistently shown to have preferential activation for faces (Pitcher et al., 2011). Examining the time locked virtual evoked response from the OFA grid points showed the same sensor space significance and waveform (Figure 2D).

### Seeded Connectivity

We predicted that the connectivity of the essential nodes would be higher for the respective reported percept. Analysis thus looked at the connectivity of the two separable occipital regions showing significance in the source reconstruction (LO and OFA, Figure 3A) separated for the reported perceptual response (vase & faces). Taking the window of the significant prestimulus low oscillatory power effect we show the iCOH for the contrasts. There was a significant main effect of seed between the responses (p < .05, *corrected*), with LO seed showing stronger overall connectivity. Finally, there was an interaction between the seed and the reported response (p < .05, *corrected*). That is, for face report there was stronger prestimulus connectivity for the OFA seed, and weaker connectivity for the LO seed. Conversely for the vase report there was stronger connectivity for the LO seed, and weaker connectivity for the OFA seed. Both effects were maximal at 7 Hz. The right inferior frontal cortex was the largest locus of the main effect and the interaction (See Figure 3).

**Figure 3:**
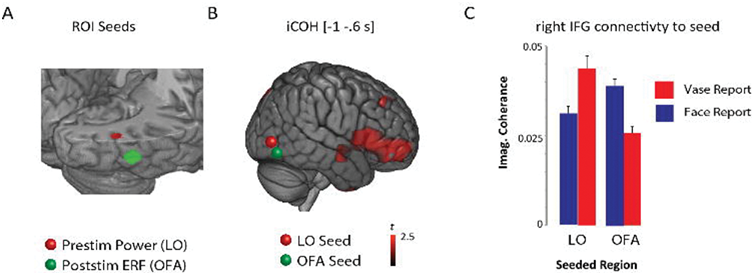
A: The two peak effects for the prestimulus (12 – 17Hz) effect, these regions were used for the ROI based seeded iCOH analysis. B: The interaction of seeds × stimulus report, the source is the masked effect (p < .05, *corrected*). C: Shows the interaction effect from the extracted grid points in frontal regions that showed significance as seen in panel B. The values presented are the absolute mean and standard deviation of the iCOH values.

## Discussion

We investigated whether prestimulus alpha/low-beta oscillatory brain states influence what we perceive when exposed to an ambiguous picture. Using the face-vase stimulus we show that alpha/low-beta power is relatively increased in the right LO before face reports. This is coherent with the theory that in situations of high rivalry, features or areas are inhibited to increase processing capacities for relevant features (Jensen and Mazaheri, 2010). The inhibition of LO reduces the probability of perceiving the vase, thus increasing the likelihood of the upcoming image to be perceived as faces.

Evoked response differences show a greater response to faces vs. vase in a late 350msec time window, and this effect is localised in the right OFA, a region with strong response to faces (Pitcher et al., 2011). Finally, we show for the first time that the state of connectivity between canonical “object” and “face “regions to inferior frontal cortex biases the outcome of the perceptual competition between Vase and Face. It corroborates a central assumption of the Win2Con framework (Ruhnau et al., 2014; Weisz et al., 2014), that: prestimulus network integration of relevant sensory modules form pre-established pathways along which neural activation can propagate, thereby shaping perceptual states.

We interpret the increase of low-frequency power in the right LO as inhibition of interfering brain activity within an area critical for object perception. Increased low-frequency oscillatory activity is associated with the inhibition of the respective brain region resulting in decreased stimulus processing (Haegens et al., 2011; Jensen and Mazaheri, 2010; Keil et al., 2014). Most of these studies showed inhibitory effects in the alpha frequency band (8–12 Hz) in the visual modality, however, depending on the brain region the exact frequency band can differ from the classical alpha band, while still having an inhibitory function (Bernasconi et al., 2011; Chen et al., 2008; Hari and Salmelin, 1997; Keil et al., 2014; Siegel et al., 2008). The 12-17 Hz effects we revealed are thus nicely in line with the high alpha/low beta effects reported in the literature and so can be used to describe an inhibitory role in stimulus processing.

The LO, the region exhibiting the main low-frequency power increase in the present study, is a composite of the LOC, as such it is one of the main modules of object processing (Grill-Spector et al., 1999; Kourtzi and Kanwisher, 2001). Most of these studies report a bilateral activation of the LO, while the effect we found was lateralized to the right hemisphere. This could be due to the type of task we used. For instance, (Large et al., 2007) showed that the right LO is activated for repetitive object presentation, McAuliffe & Knowlton (2001) implicated the right LO as being specialized for object identification, while Vuilleumier, Henson, Driver, & Dolan (2002) postulated that the right LO has a higher sensitivity to object view. The source of the power effect is associated with an area known to be a key hub of object processing (LO) and not to an area critical in face processing (OFA), which is in close proximity. This suggests that alpha/low-beta power is indeed enhanced in the region preferentially processing objects, pointing to an inhibition of that area before participants see the faces.

Ongoing oscillations could indeed be spontaneous in design and the effects observed could derive from a purely random fluctuation in brain activity in line with the interpretation of Hess Elman et al. (2008) fMRI results within the face-vase illusion. Another interpretation is top-down preparatory response, whereby participants decided what to see before they saw it. These responses have been shown to be in object processing sources (Peelen and Kastner, 2011) rather than V1 and indeed match up with the results we observe, and are a possible explanation for these effects. This also, along with a possible motor preparation effect could be a reason why we observe the starting bias within the DDM analysis (Bode et al., 2012), furthermore inspection of the post stimulus time frequency period revealed a topography in the beta frequency range which would adhere to a difference between the motor preparation between the two reports (see Inline Supplementary Figure 3). Within the current study, which was intentionally kept very close to the original fMRI study by Hesselmann et al. (2008), this issue cannot be resolved with certainty and requires further studies. The trial-by-trial switching that occurred was at random, although previous research has shown that trial-by-trial reports of rivalrous stimulus material is stable if the stimulus is disrupted during a dominant percept (Leopold et al., 2002). However, it is hypothesized that one of the key reasons behind this stabilization is due to a peak dominant percept effect (Wilson, 2007) and when a percept is not full dominant or indeed is presented briefly then the stabilization could be lost. Within our presentation parameters we *destabilized* the dominance three-fold by having presentation brief, masked and a long jittered inter trial period.

The brief and late latency effects we observed in the event-related fields suggest that there were some processing differences that altered the report of face vs. vase. Although generated from a known face sensitive area OFA, the specifics are harder to disentangle. We did not observe differences within the M170 (or ERP N170) component commonly observed within the range of 120-200msec peaking around 170 msec after stimulus onset (Halgren et al., 2000), however previous research with the face-vase stimulus has shown late effects (Pitts et al., 2011). Even though evoked responses were descriptively enhanced for face reports in this more classical time-interval, the difference was statistically not significant. This maybe due to the approach we took in analysing the data. Researchers looking at M170 effects often select channels on the basis of an individual effect derived from a localiser (Xu et al., 2005). Indeed fMRI seems to yield a large FFA response when attending to the face within the face-vase stimuli (Andrews et al., 2002). There are however other studies which show lack of attenuation of the M170 in scene clutter (Andalman and Sinha, 2010) which suggests that the effects occur before segmentation of the face-vase stimulus. Thus, the absence of an M170 effect is not surprising given that the segmentation always occurs later than the M170. A possible explanation is that we observed a later face-selective process, modulated by attention. This is suggested by the source analysis, which was localised in the OFA an area critical for face processing (Steeves et al., 2006), which has been shown to be modulated by attention (Haxby et al., 2000).

Our studies main goal was to describe the network integration with respect to the upcoming perceptual report. This analysis is of crucial importance within the context of our framework, in which we state that prestimulus local excitability states are only playing a minor role in driving upcoming percepts (Ruhnau et al., 2014). We proposed that relevant sensory processing modules (i.e. essential nodes) require pre-established connections to high-level (frontal/parietal) areas to enable upcoming conscious perception. In cases of ambiguous stimuli, we predict that specific prestimulus coupling and decoupling patterns of category sensitive regions to frontal and parietal regions can bias perceptual competition towards one percept or the other. The present study fully confirms this prediction: Local canonical areas of object and face processing (LO & OFA) show a strong coupling to right inferior frontal cortex - a putative region of global access to consciousness (Dehaene and Changeux, 2011) - prior to the respective perceptual reports. The network state is seen already prior to stimulus onset, suggesting that ongoing network states constitute "pre-established pathways of information flow" when confronted with sensory stimuli (Weisz et al., 2014). However, the framework is also refined for the analysis of conscious content and the contrastive constraints of having a larger stimulus feature space required us to also look for essential nodes present in the evoked time window.

This is the first time that prestimulus connectivity has been used in a bistable paradigm to assess the idea that connectivity states represent relevant prerequisites to consciousness (Aru et al., 2012). Our study corroborates findings from more often used near-threshold or masking paradigms. Using an ambiguous stimulus comes with a crucial extension to the purely ‘seen vs. unseen’ type of task: participants always reported either a vase or faces, thus a conscious "content" was always present. Our results a demonstration that in cases of supra-threshold presentations, specific network states preceding the stimulation shape the content of upcoming perception. This further ratifies the previous fMRI studies, and how the prestimulus BOLD can highlight category content prior to stimulus processing (Hesselmann et al., 2008). Indeed, MEG and its strength of large-scale dynamic connectivity analysis are, within this study, able to precisely track the unfolding of prestimulus brain states.

In the present study we exposed subjects with ambiguous stimuli, here the face-vase stimulus, to investigate determinants of the content of consciousness in oscillatory brain dynamics. We show that the local modular effects are coupled with large-scale interareal connectivity. This supports the notion of the Win2Con framework that it is not merely the local prestimulus states that can shape what is the content of consciousness but open windowed/connected brain states. Furthermore, we show a frequency effect that spans two frequency bands, this further suggests that the rigid ideas of frequency band specific effects and their ties to specific cognitive mechanisms should be applied more liberally as frequency bands may share more common properties.

### Acknowledgments

NP, NM, PR, and NW were supported by the European Research Council ERC StG 283404 (WIN2CON). We thank Marius Peelen and Thomas Hartmann for constructive feedback in early stages of the manuscript preparation.

